## Supporting Information for "The AMP deaminase of the mollusk *Helix pomatia* is an unexpected member of the adenosine deaminase-related growth factor (ADGF) family"

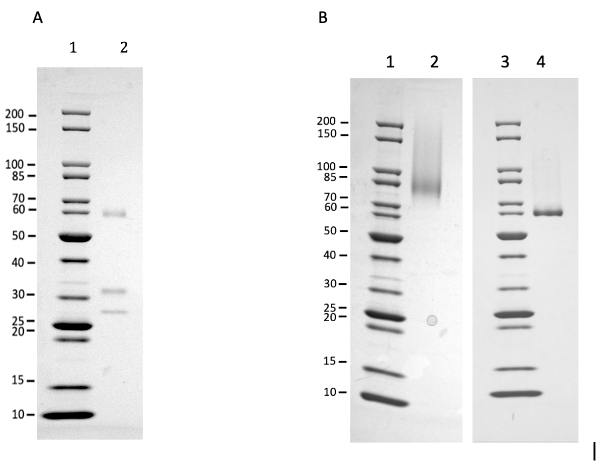

**S1 Fig**. **Polyacrylamide SDS gel electrophoretogram of native and recombinant HPAMPD.** Panel A. Lane 1 is MW standard. Lane 2 is native HPAMPD.

Panel B. Lane 1 and 3 are MW standards. Lane 2 is HPAMPD prior to deglycosylation with EndoH_f_. Lane 4 is HPAMPD after EndoH_f_ treatment.

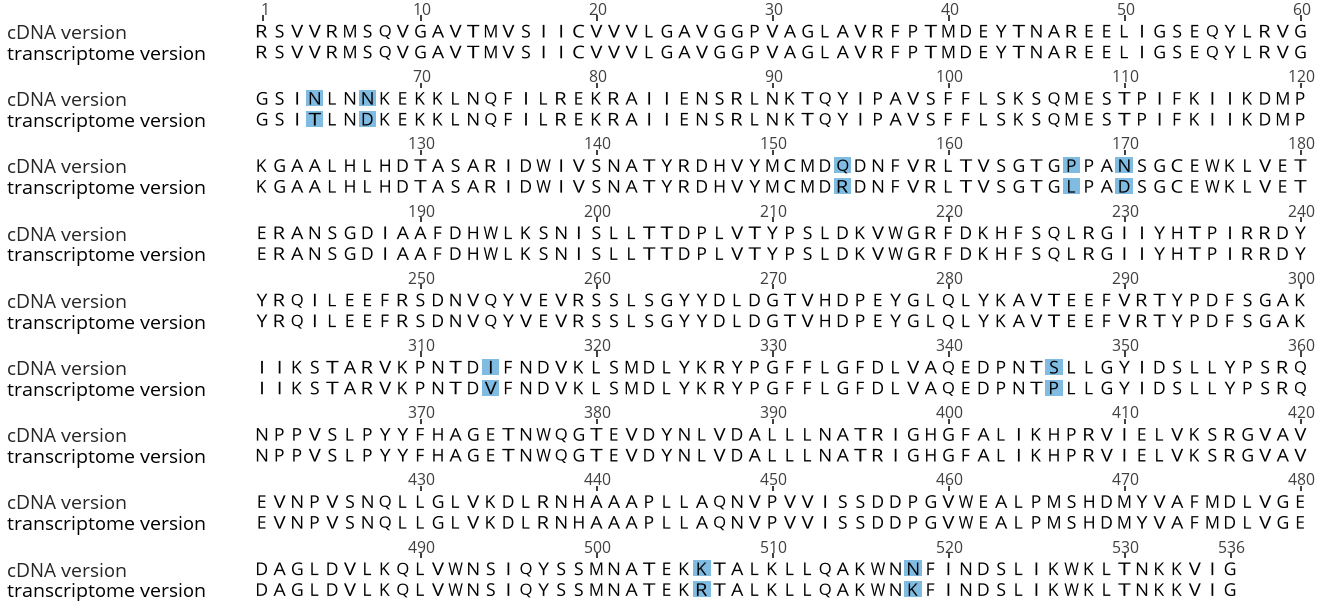

**S2 Fig. An alignment of the amino acid sequence of the translated cDNA with the translated contig 71391 from the *H. pomatia* transcriptome.** The amino acid differences are highlighted in blue.

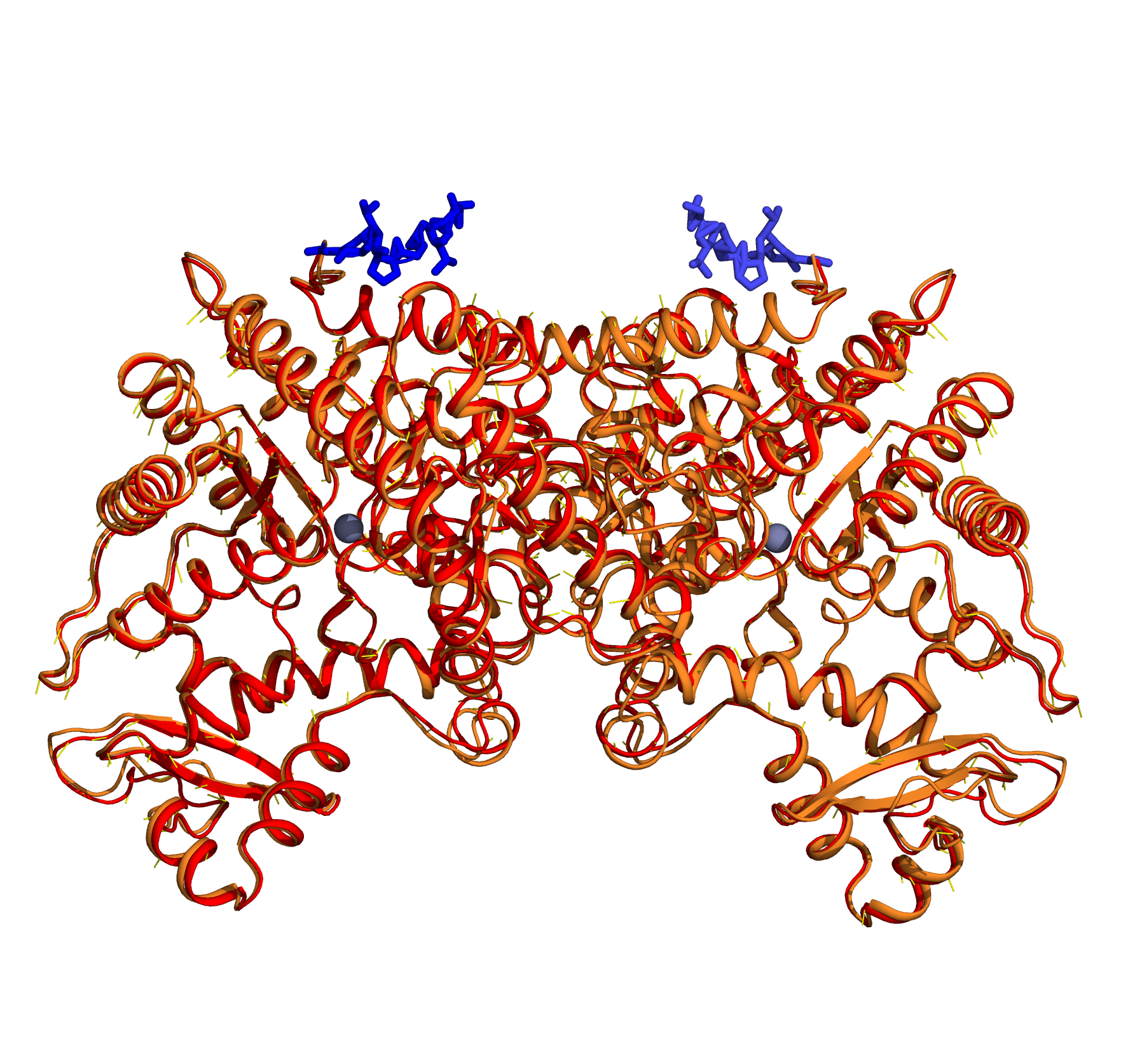

**S3 Fig**. Superimposition of HPAMPD homodimers that were predicted with AlphaFold (Jumper, 2021) from amino acid sequences with and without the additional 8 terminal residues AVGGPVAG (blue sticks). The gold cartoon was predicted from sequence with the additional 8 residues, the red cartoon was predicted from sequence without the extra residues.

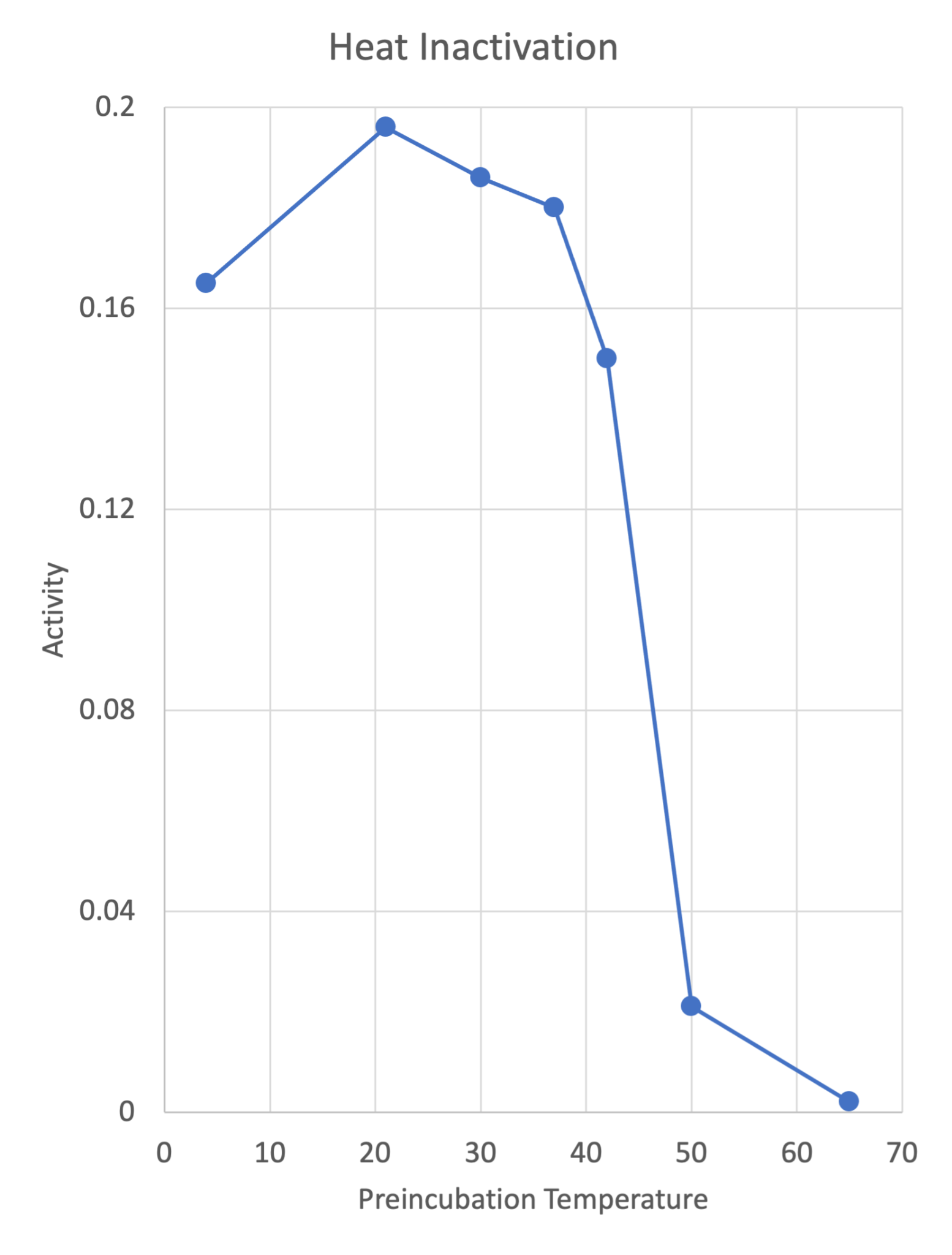

**S4 Fig.** **HPAMPD heat inactivation**. The HPAMPD was incubated in 10 mM Tris-HCl pH 7.5, 1 mMEDTA, 50 mM NaCl for 30' at either 4°, 20°, 30°, 42°, 50° or 65° and then assayed for activity.

**S1 Table. List of identified peptides from MS analyses of the digests (Peptides_identified_protein.xlsx).**

**S2 Table. List of correspondence of Transcriptome Contig numbers to Protein IDs**

**(Transcriptome correspondence.xlsx)**
